# The Biogeography of *Dromiciops* in Southern South America: middle Miocene transgressions, speciation and associations with *Nothofagus*

**DOI:** 10.1101/2020.08.12.207332

**Authors:** Julian F. Quintero-Galvis, Pablo Saenz-Agudelo, Juan L. Celis-Diez, Guillermo C. Amico, Soledad Vazquez, Aaron B.A. Shafer, Roberto F. Nespolo

## Abstract

**Aim:** Several geological events affecting Southern South America during the middle Miocene climatic optimum acted as important drivers of diversification to the biota. This is the case of Microbiotheria, for which *Dromiciops* is considered the sole surviving lineage, the sister group of Eomarsupialia (Australian marsupials). Three main *Dromiciops* genetic lineages are known, whose divergence was initially attributed to recent Pleistocene glaciations. Using fossil-calibrated dating on nuclear and mitochondrial genes, here we reevaluate this hypothesis and report an older (Miocenic) biogeographic history for the genus.

**Location:** Southern South America.

**Methods:** Phylogenetic reconstruction using sequences from two mitochondrial DNA and four nuclear DNA genes in 159 specimens, from 31 sites across Chile and Argentina. Divergence time estimation using fossil calibration.

**Results:** Our phylogenetic analysis resolved four well supported clades with discrete geographic distributions. The oldest and most differentiated clade corresponds to that of the northern distribution (35.2°S to 39.3°S), which would be a different species (*D. bozinovici*, sensu D’elia et al. 2016). According to our estimations, this species shared a common ancestor with *D. gliroides* (southern clades) about 13 million years ago (95% CI: 6.4-25.3). The southern clades (39.6°S to 42.0°S), showed a divergence time ranging from 9.57 to 6.5 Mya. Strong genetic structure was detected from north to south but not across the Andes, or between Chiloé island/ mainland. Demographic equilibrium is inferred to the northern clade, and recent demographic expansions was detected in the central and southern clades.

**Main conclusions:** The whole diversification of *Dromiciops* occurred within the Miocene, being the Middle Miocene transgression (MMT), the massive marine flooding that covered several lowlands of the western face of los Andes between 38-48° S, the most likely diversifying force. This was the result of an increase in global sea levels due to the Miocene climatic optimum, which shaped the biogeographic origin of several species, including *Nothofagus* forests, the habitat main of *Dromiciops*.

## Introduction

Several forces have been associated to the mammalian radiation that was initiated in the Cenozoic, which in South America was marked by the disconnection of the Antarctic bridge in the Eocene (Hershkovitz, 1999). Since then, the Southern South American mammalian fauna suffered a number of climatic constraints, fragmentations and redistributions that ended in a faunal assemblage with high degree of endemism (Habel et al., 2010). This is the case of the marsupial “Monito del monte” (genus *Dromiciops* Thomas, 1894), which represents one of the most intriguing cases of relict species (sensu Habel *et al.*, 2010). It is the only extant species of the order Microbiotheria, the sister group of Eomarsupialia, otherwise known as Australian marsupials (Hershkovitz, 1999; Nilsson *et al.*, 2004; Mitchell *et al.*, 2014). Fossil evidence suggests that Microbiotheria was originated around 49Mya and diversified in the middle Eocene; most species subsequently disappeared from the fossil record by the end of the early Miocene, except *Dromiciops* (Marshall & de Muizon, 1988; Goin *et al.*, 2010; Goin & Abello, 2013). Microbiotheria distributional range likely suffered a strong constriction after the orogenesis of Los Andes, in the Miocene, due to a reduction in the extension of the temperate rainforest of South America, the main habitat of microbiotheriids (Abraham *et al.*, 2000; Garzione *et al.*, 2008).

Today, *Dromiciops* is restricted to the Valdivian temperate rainforest (“Valdivian rainforest”, hereafter) (Martin, 2010), from the Maule Region (35°50’S, 72°30’W) to Chaitén (45°00’S, 72°00’W) in Chile (Oda *et al.*, 2019), and from Neuquén Province to the Río Negro Province in Argentina (Martin, 2010; Celis-Diez *et al.*, 2012; Gurovich *et al.*, 2015). Although several authors have explored distinct genetic and taxonomic aspects of this group (Himes *et al.*, 2008; D’Elía *et al.*, 2016; Valladares-Gómez *et al.*, 2017, 2019; Martin, 2018), several aspects of its evolutionary history and genetic structure have yet to be fully resolved.

Himes *et al.* (2008) evaluated *Dromiciops* genetic structure using two mitochondrial fragments and observed the existence of three well-differentiated clades and a marked geographic structure along the sampled range comprising Chile and Argentina. These authors concluded that the southern clade probably experienced a recent expansion, while the northern clade exhibited long-term stability. Despite reporting around 10% genetic divergence among clades, Himes et al. (2008) attributed this divergence to inter-population variation and recent climatic phenomena (i.e., Pleistocene glaciations). This view was challenged by D’Elía *et al.* (2016) that reinterpreted the genetic evidence and added morphometric analyses, and proposed two new *Dromiciops* species; *D. bozinovici* (northern clade) and *D. mondaca* (coastal area near Valdivia) with *D. gliroides* being relegated to the southern distribution. Additional morphological and genetic structure analyses have been performed (Valladares-Gómez *et al.*, 2017; Martin, 2018; Suárez-Villota *et al.*, 2018), but accurate estimations of divergence times and detailed genetic analyses at the level of the whole geography have yet to be conducted.

Here we provide a fossil-calibrated divergence time obtained from a panel of genetic markers, for the major *Dromiciops* lineages of Chile and Argentina. This information permitted us to test explicit biogeographic hypotheses for the origin and diversification of this endemic marsupial, together with an assessment of population connectivity. Specifically, we tested whether *Dromiciops* lineages are explained by the last glaciations which fragmented the southern populations (i.e. a “Pleistocene” scenario) as originally proposed by Himes et al. (2008). Alternatively, we considered an older diversification scenario involving geological events occurring in the Middle Miocene, coincident with the biogeographic history of *Nothofagus* forests (Acosta *et al.*, 2014).

## Material and Methods

### Sample collection, DNA extraction, and sequencing

Between 2008 and 2019, samples were collected from 13 localities across Chile and Argentina. In most cases, animals were live captured using tomahawk traps installed during nighttime at each locality in summer; in a few cases samples correspond to animals found dead but in good condition and stored frozen at -20°C. When captured alive, a small skin clip was taken from the ear of each individual and it was then immediately released into the same place where it was captured. The entire procedure was performed according to the bio-ethical guidelines of the Universidad Austral de Chile (UACh Project license number 70/8818 and 313/2018) and SAG wildlife (Project license number 70/8818, 4371/2019 and 3393/2019, held by Roberto F. Nespolo). Tissue origin, locations and sample sizes are provided in Table 1 and Fig 1. Tissues were stored in 96% ethanol and DNA was extracted using GenJet Genomic DNA (Thermo Fisher Scientific).

**Table 1.** Information of the localities and samples used in this study. Codes in column markers available are: a (Cytb), b(12sRNA), c(vWF), d(IRBP), e(ApoB), f(RAG-1).

| Code | Locality name | Number of samples | Region or Province | Country | Latitude | Longitude | Markers available | Paper description |
| --- | --- | --- | --- | --- | --- | --- | --- | --- |
| 1 | Ñuble, San Fabian | 1 | Bio Bio (VII) | Chile | -36.5833 | -71.4667 | a,b,c,d,e,f | Himes et al. 2008 |
| 2 | Caramavida | 7 | Bio Bio (VII) | Chile | -37.6960 | -73.1969 | a,b,c,d,e,f | This study |
| 3 | Nahuelbuta | 11 | Araucanía (IX) | Chile | -37.8245 | -71.1833 | a,b,c,d,e,f | This study |
| 4 | Malleco, Angol, Vegas Blancas | 2 | Araucanía (IX) | Chile | -37.8333 | -72.8667 | a,b,c,d,e,f | Himes et al. 2008 |
| 5 | Malleco, Curacautín, Termas de Tolhuaca | 3 | Araucanía (IX) | Chile | -37.8245 | -72.9712 | a,b,c,d,e,f | Himes et al. 2008 |
| 6 | Cajón de Aguas Negras | 1 | Araucanía (IX) | Chile | -38.5333 | -71.1833 | a,b,c,d,e,f | Himes et al. 2008 |
| 7 | Cautín, Cunco, East of Lago Colico | 1 | Araucanía (IX) | Chile | -39.1000 | -71.8667 | a,b,c,d,e,f | Himes et al. 2008 |
| 8 | Aluminé, Quillén | 2 | Neuquén | Argentina | -39.3678 | -71.2339 | a,b,c,d,e,f | Suarez-Villota et al. 2018 |
| 9 | Aluminé, Lago Quillén | 4 | Neuquén | Argentina | -39.3667 | -71.2333 | a,b,c,d,e,f | Himes et al. 2008 |
| 10 | Fdo. San Martín 1 | 11 | Los Ríos (XIV) | Chile | -39.6288 | -73.1870 | b,c,d,e,f | Himes et al. 2008; Suarez-Villota et al. 2018 |
| 11 | Fdo. San Martin 2 | 11 | Los Ríos (XIV) | Chile | -39.6500 | -73.1941 | a,b,c,d,e,f | This study |
| 12 | Parque Oncol | 12 | Los Ríos (XIV) | Chile | -39.7027 | -73.3263 | a,b,c,d,e,f | This study |
| 13 | Fdo Los Guindos | 1 | Los Ríos (XIV) | Chile | -39.6288 | -73.1870 | a,b,c,d,e,f | Himes et al. 2008 |
| 14 | Forestal Calle Calle 1 | 13 | Los Ríos (XIV) | Chile | -39.8667 | -73.0500 | a,b,c,d,e,f | This study |
| 15 | Forestal Calle<br>Calle 2 | 4 | Los Ríos (XIV) | Chile | -39.8667 | -73.0500 | b,c,d,e,f | Suarez-Villota et al. 2018 |
| 16 | Fdo. Senderos del<br>Bosque | 1 | Los Ríos (XIV) | Chile | -39.6288 | -73.1870 | a,b,c,d,e,f | Suarez-Villota et al. 2018 |
| 17 | Parque Llancahue | 10 | Los Ríos (XIV) | Chile | -39.8596 | -73.1373 | a,b,c,d,e,f | This study |
| 18 | Donoso | 6 | Los Ríos (XIV) | Chile | -39.8658 | -73.1541 | a,b,c,d,e,f | This study |
| 19 | Entre Lagos | 2 | Los Lagos (X) | Chile | -40.6667 | -72.2667 | a,b,c,d,e,f | Himes et al. 2008 |
| 20 | Villa La Angostura | 2 | Neuquén | Argentina | -40.6675 | -71.7042 | b,c,d,e,f | Suarez-Villota et al. 2018 |
| 21 | Los Lagos,<br>Estancia Paso<br>Coihue | 1 | Río Negro | Argentina | -40.8833 | -71.2833 | a,b,e,f | Himes et al. 2008 |
| 22 | Isla victoria | 1 | Neuquén | Argentina | -40.8967 | -71.5417 | b,c,d,e,f | This study |
| 23 | Puerto Blest | 5 | Río Negro | Argentina | -41.3574 | -71.5103 | a,b,c,d,e,f | This study |
| 24 | Llao-Llao<br>Bariloche | 7 | Río Negro | Argentina | -41.0333 | -71.5333 | a,b,c,d,e,f | This study |
| 25 | La Picada | 3 | Los Lagos (X) | Chile | -41.3574 | -71.5103 | a,b,c,d,e,f | Himes et al. 2008 |
| 26 | Llanquihue, East<br>of Lago<br>Llanquihue | 1 | Los Lagos (X) | Chile | -41.1833 | -72.5500 | a,b,c,d,e,f | Himes et al. 2008 |
| 27 | Laguna Verde | 14 | Río Negro | Argentina | -41.3574 | -71.5103 | a,b,c,d,e,f | This study |
| 28 | Peninsula Llao-Llao, Bariloche | 2 | Río Negro | Argentina | -41.5197 | -71.5625 | a,b,c,d,e,f | Himes et al. 2008 |
| 29 | Caulin Chiloe | 7 | Los Lagos (X) | Chile | -41.8333 | -73.6090 | a,b,c,d,e,f | This study |
| 30 | Senda Darwin<br>Chiloe | 10 | Los Lagos (X) | Chile | -41.8691 | -73.6830 | a,b,c,d,e,f | This study |
| 31 | Ancud, Chiloé | 3 | Los Lagos (X) | Chile | -41.8833 | -73.6833 | a,b,c,d,e,f | Himes et al. 2008 |

**Figure 1.**
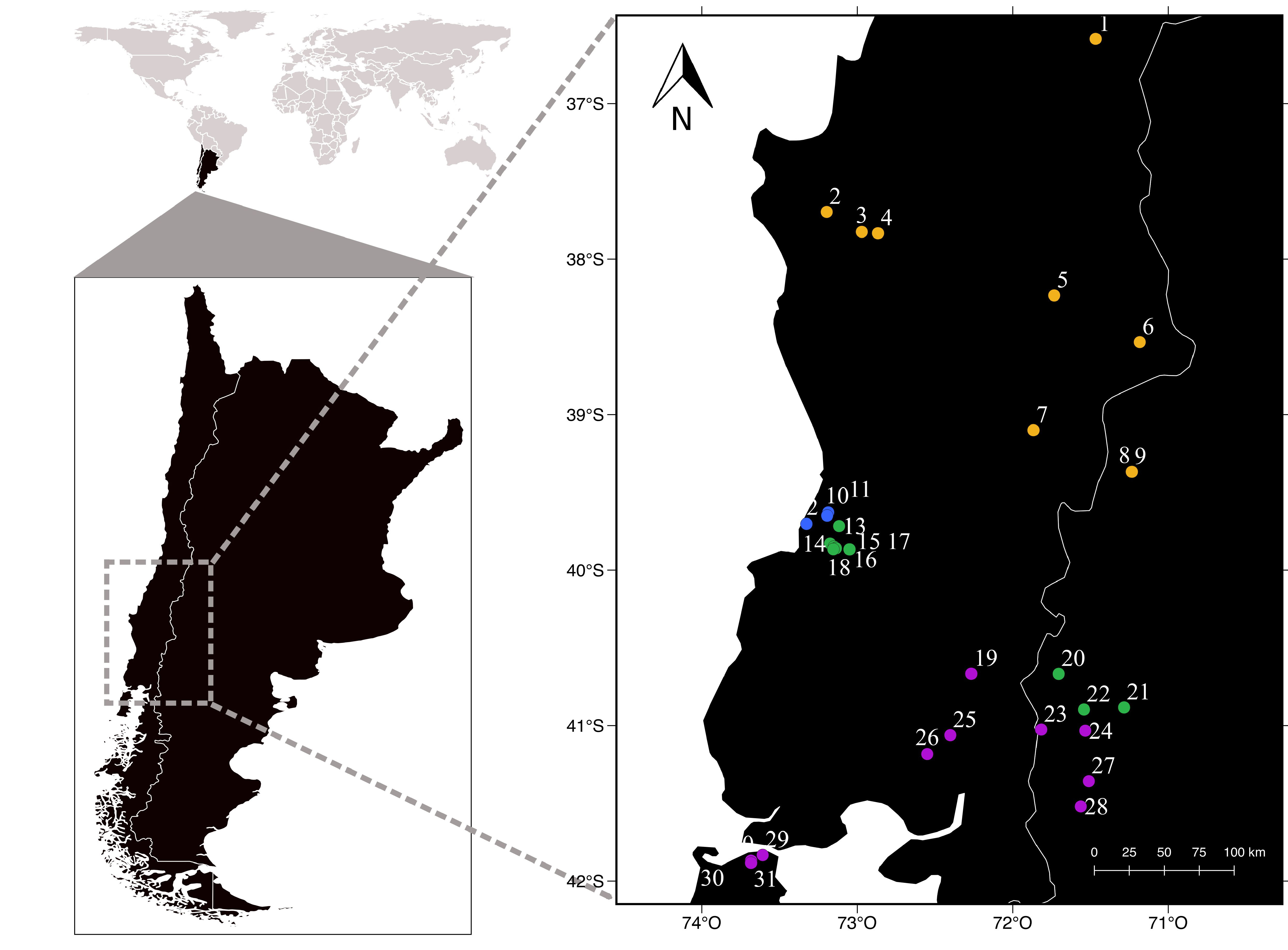
Sampling sites across Argentina and Chile. Numbers correspond to those of Table 1. Colors point correspond to the phylogeny clades in figure 2.

For all sampled individuals, we sequenced two mitochondrial genes (12S RNA and cytochrome b [Cytb]) (Smith & Patton, 1993; Palma & Spotorno, 1999; Himes *et al.*, 2008) and four nuclear genes (apolipoprotein B gene [ApoB], recombination activating gene-1 [Rag-1], interphotoreceptor retinoid binding protein gene [IRBP] and von Willebrand factor gene [vWF]) (Stanhope *et al.*, 1992; Springer *et al.*, 1997; Amrine-Madsen *et al.*, 2003). PCR reactions were carried in 10μL that included 1µl of genomic DNA, 3.5 µl of GoTaq Colorless Master Mix (Promega), 0.6µl of forward and reverse primers, 0.2 BSA and 4.1 of ultrapure water. Cycling conditions for all assays were as follows: initial denaturation at 94°C for 10 min; 35 cycles at 94°C for 1 min (denaturation), 40°C (Cytb)/55°C (all other genes) for 2 min (annealing), and 72°C for 2 min (elongation); and final elongation at 72°C for 13 min. We used the software Geneious v.11.1.4 (Kearse *et al.*, 2012) for filtering, trimming and alignment of sequences. We also complemented our sampling scheme by adding 45 sequences of *D. gliroides* of four nuclear genes and 12S RNA from 18 localities previously published by Suárez-Villota et al. (2018) and 31 sequences of Cytb sequences from Himes *et al.* (2008).

All coding sequences were translated into amino acids to corroborate the absence of internal stop codons that could indicate the presence of pseudo-genes. Sequences were aligned using MAFFT (Katoh & Standley, 2013) as implemented in Geneious with default parameters. For the nuclear markers, some samples were polymorphic at multiple positions and gametic phases were ambiguous. Therefore, all sequences were checked in the forward– reverse nucleotide assemblies to ensure that all double peaks were correctly identified using standard degeneracy codes. We then inferred the gametic phase using DnaSP v5 (Librado & Rozas, 2009).

### Phylogenetic analysis

We reconstructed *Dromiciops* phylogenetic relationships using mitochondrial and nuclear markers and Maximum Likelihood (ML) and Bayesian Inference (BI). These analyses were performed two times: once concatenating mitochondrial markers and nuclear markers in two separate data sets, and once as a fully concatenated dataset (nuclear + mitochondrial markers). ML phylogenetic analysis were performed in IQ-TREE v1.6.8 (Nguyen *et al.*, 2015), with 1000 bootstrap replicates, using default setting. BI trees were implemented in MrBayes v3.2.1 (Ronquist *et al.*, 2012). The most appropriate nucleotide substitution models for every gene alignment and concatenated segments were selected using the Akaike Information Criterion as implemented in JMODELTEST 2.1.4 software (Darriba *et al.*, 2012). Posterior distributions of BI phylogenies were obtained by a Markov Chain Monte Carlo (MCMC). Samples of the trees and parameters were drawn every 100 steps from a total of 3 × 10^7^ MCMC generations. Three additional runs were conducted beginning with random trees. The 50% majority rule consensus of the post-burn (using a burn-in of 25%) for all of the generations was computed across all runs. Trees were visualized and edited in FIGTREE 1.4 (Rambaut & Drummond, 2012). We built Media-Joining haplotype networks for each of the genes and mtDNA and nDNA concatenated separately. The program POPART (Leigh & Bryant, 2015) was used to build a Median-Joining (MJ) network for both mtDNA and nDNA data to reconstructed a haplotype network.

### Species delimitation

We applied two multi-locus coalescent-based methods for delimiting potential new *Dromiciops* species. First, we used a Bayes factor delimitation (BFD) approach (Grummer *et al.*, 2014), to compare alternative species trees and to determine the number of diagnosable evolutionary lineages in *Dromiciops.* Second, we used the main clades identified in the phylogenetic analyses (see results) to test species delimitation hypotheses with Bayesian species delimitation analysis implemented in the program BPP version 3.3 (Yang, 2015). This program uses the multispecies coalescent model to compare posterior probabilities assigned to the major phylogenetic groups (Yang & Rannala, 2010). We performed separate A10 (species delimitation on a guide tree) and A11 (joint species delimitation and species tree estimation) analyses using empirical priors calculated from the data. We also measured the genealogical divergence index (*gdi*) among clades (Jackson *et al.*, 2017; Leaché *et al.*, 2020) for evaluating where populations lie on the speciation continuum. This index compares genetic divergences among population pairs of different species in a coalescent framework that considers the joint effects of genetic drift and migration. We implemented the A00 output of BPP (estimation of parameters θ and τ, under a multispecies coalescent model) to calculate the *gdi* following the equation: *gdi* =1–e^-2τ/ θ^ (Jackson *et al.*, 2017). As suggested by Jackson et al, (2017), clades or populations can be considered distinct species if *gdi* values are > 0.7, while *gdi* values below 0.2 indicate populations belonging to one single species.

### Estimation of divergence time

Times of divergence were estimated using a strict clock Bayesian approach in BEAST v2.5 (Drummond *et al.*, 2012). We used the substitution model suggested by JMODELTEST 2.1.4 software (Darriba *et al.*, 2012) (Table S2), including a coalescent constant population model selected as prior, and a random starting tree. We calibrated and constrained the internal age of our nodes based on Didelphimorphia and Eomarsupialia as outgrups, using 14 fossil records (Table S3 in Mitchell *et al.*, 2014) distributed across the marsupial phylogeny and assuming a uniform calibration priors (Drummond *et al.*, 2012). Three independent MCMC runs were performed with 3 × 10^7^ generations sampled every 5000 steps with 10% samples discarded as burn-in, and the output was tested in Tracer v1.7.1 (Rambaut *et al.*, 2018). The effective sample size (ESS) of each parameter was required to be higher than 200. The final tree was summarized using the program TreeAnnotator included in BEAST v2.5.

### Molecular diversity and genetic structure

Standard genetic diversity indices, such as the number of haplotypes (H), polymorphic sites, haplotype diversity (H_D_), and nucleotide diversity (π), were calculated using DnaSP v5 (Librado & Rozas, 2009). The software DnaSP was also used for estimating the average number of nucleotide substitutions per site between populations (D_xy_) (Nei, 1987) and average number of nucleotide differences between populations (K_xy_) (Tajima, 1989). An analysis of molecular variance (AMOVA) (Excoffier *et al.*, 1992) was executed to investigate the genetic structure of populations using the Arlequin software (v3.5) (Excoffier & Lischer, 2010). Distinct hierarchical analyses of genetic diversity were performed, which differed in the way sampling locations were grouped into clusters. Clusters were defined based on the resulting phylogenetic tree and haplotype networks. For every gene, we applied two different models. Model I (2 groups): clade A vs (clade B + C + D); Model II (3 groups): clade A vs clade B vs clade C-D. Genetic differentiation and genetic distances among populations were estimated by calculating the Phi-statistic (Φ_ST_) between populations and testing their significance with 1000 permutations using Arlequin.

Isolation by distance (IBD) for concatenated mitochondrial and nuclear markers were analyzed in GenALEX v6.5 (Peakall & Smouse, 2012) to test for a correlation between genetic (Φ_ST_) and geographic distance matrices. We defined geographic distance as the linear distance between every population pair, and population pairwise Φ_ST_ as genetic distances estimated in Arlequin v3.5 (Excoffier & Lischer, 2010). The correlation between the two distance matrices was assessed using a Mantel test, with the significance estimated after 100.000 random permutations.

### Historical demography

We used a variety of approaches to assess the demographic history of each clade. First, we used two neutrality statistics, Tajima’s D and Fu’s Fs calculated with 10,000 random permutations using Arlequin v3.5 (Excoffier & Lischer, 2010) to test for population expansion events and deviations from a strictly neutral model of evolution. This was done in 16 localities with samples sizes larger than four individuals. Negative values for these indices are indicative of either population expansion or natural selection (Tajima, 1989; Fu, 1996). We calculated the pairwise mismatch distribution per clade and global using Cytb gene (Rogers & Harpending, 1992), and compared it to the expected distribution under a population growth-decline model in DnaSP.

## Results

### Sequence data

After trimming and alignment, we recovered 417 base pairs (bp) for Cytb, 764 bp for 12S RNA, 708 bp for ApoB, 951 bp for IRBP, 516 bp for RAG, and 948 bp for vWF. The best substitution models were the same for mitochondrial genes (model GTR+I+G) but differed for all nuclear genes (Table S2). This completed a total sample size of 159 individuals from 31 sites (Table S1). All new sequences were deposited in GenBank with the accession numbers -.

### Phylogenetic trees

Both the Bayesian and Likelihood phylogenetic reconstructions using the concatenated sequences (mtDNA + nDNA) recovered a similar topology with four clades (referred to clade A, B, C and D from now on Fig 2). Both the ML and IB analysis gave good statistical support for the first node separating clade A (i.e., individuals from localities 1 to 9 in Table 1; orange points in Fig 1) from samples comprising clades B-D (100/1 for ML and B respectively) (Fig 2). The support for the node separating clade B from C was moderate and differed between analyses (97/0.91 for ML and B respectively). Interestingly, clade B is restricted almost exclusively to individuals from a small area at the coastal range north of Valdivia, including the localities of San Martin (points 10 and 11 at Fig 1) and Oncol (point 12, Fig 1). The only two exceptions were two individuals (UACH 7192, UACH 7193) from a locality located across the Cruces river (Calle-Calle) less than 20 km away (Fig 2) that also grouped within this clade while all the others from the same location grouped within clade C. Further subdivisions within clade B had in general lower support (less than 80/0.5) except one node that separated samples from the localities of San Martin and Oncol sequenced in this study (99/0.81). The remaining group of samples were in turn divided into two main groups with moderate support (98/0.95) here onwards referred to as clades C and D. Clade C comprises all samples from the localities near Valdivia (Forestal Calle-Calle, Donoso y Parque Llancahue) and samples from the two localities further north of Argentina (Neuquen and Rio Negro). Clade D includes samples from Chiloe Island, and all other southern localities from Chile and Argentina (Fig 2).

**Figure 2.**
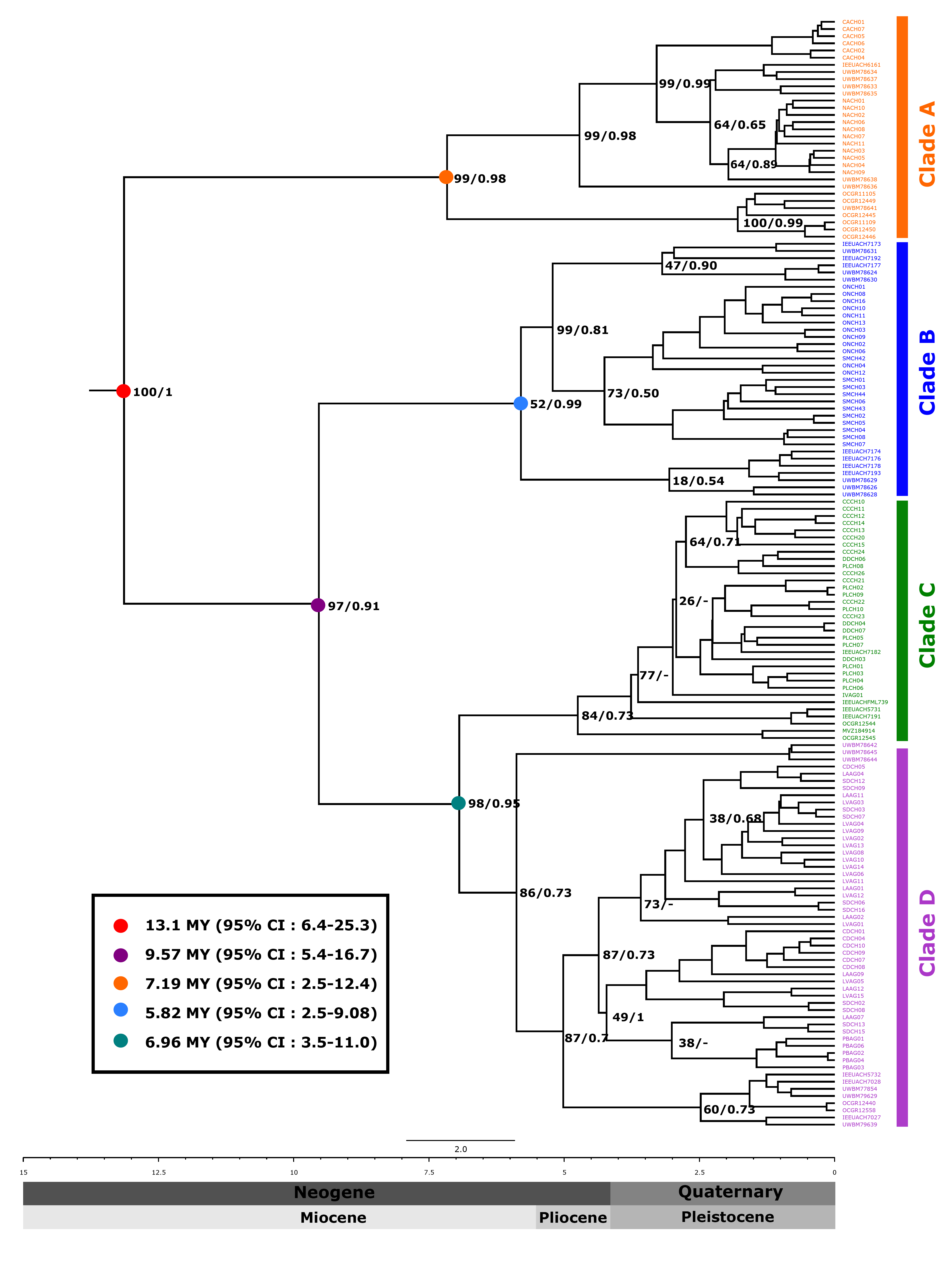
Maximum clade credibility tree inferred from IB (Cytb + 12S RNA + ApoB + IRBP + RAG1 + vWF). Values in nodes correspond the posterior probability (PP) and bootstrap (BS) support values obtained in the Bayesian inference and Maximum likelihood analysis, respectively. Color tips represent localities as assigned in each clade. The point in nodes represent assignation value for divergence times (in millions of years) for each clade, calibrated with a multilocus. Vertical bars correspond to the main clades.

### Genetic diversity within and among clades

The percentage of divergence among clades was high, ranging from 7.32 to 12% for mtDNA and 0.78 to 6.4 % for nuclear genes (Table S7). Within-clade nucleotide diversity (π) ranged between 0.007 (clade D) to 0.010 (clade A) for mtDNA, while nDNA ranged from 0.0002 (clade A for ApoB) to 0.0800 (clade C for ApoB) (Table 2). Among mtDNA, the 12S RNA gene had the highest haplotype diversity (H = 56). For the nuclear genes, ApoB was the most variable in terms of genetic diversity and haplotype diversity, while RAG-1 was the most conserved gene (Table 2 and Table S6). Pairwise comparison of the genetic differentiation indexes (Φ_ST_) for mtDNA ranged between 0.001 and 0.800 (Table S8), while nDNA ranged between 0.01 to 0.70 (Table S9).

**Table 2.**
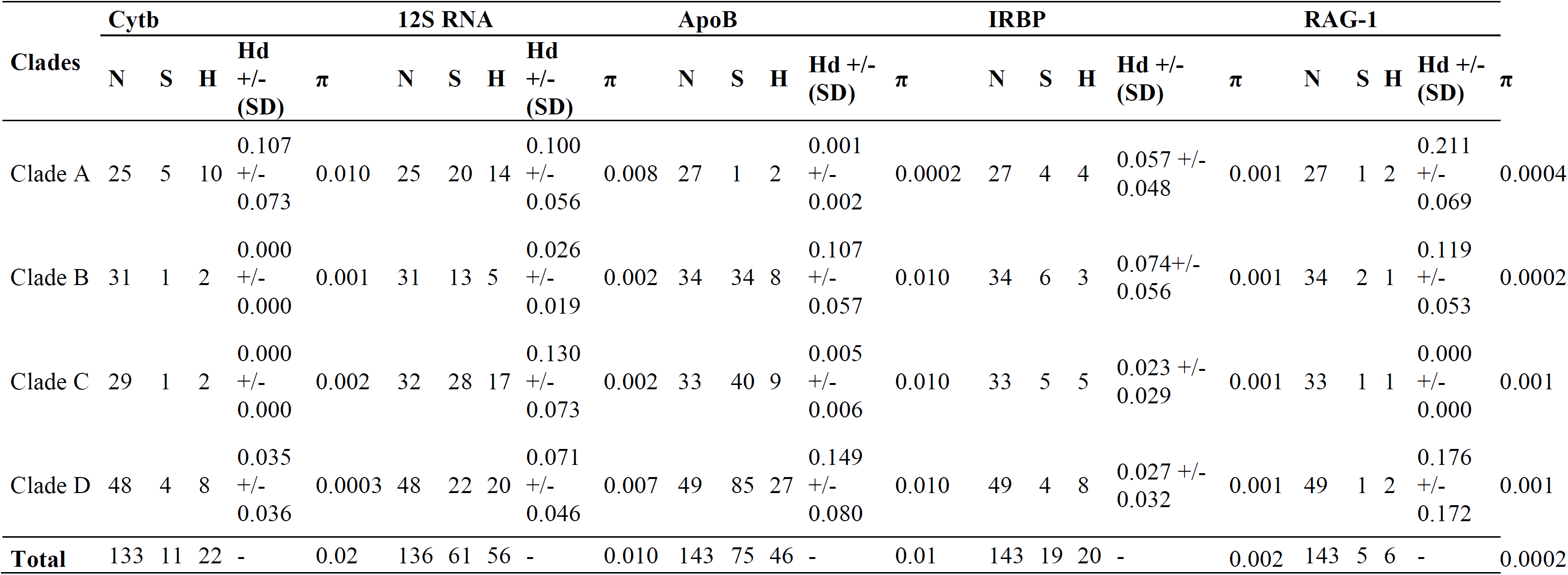
Genetic diversity indices estimated for *Dromiciops* clades. N = number of individuals; S = number of segregating sites; H = number of haplotypes; π = nucleotide diversity.

### Estimation of divergence time

Using a strict clock Bayesian approach, the estimated divergence time between clades A and (B,C,D) was 13.1 My with a 95% confidence interval of 6.4 - 25.3 My. Divergence time between B and (C,D) was estimated as 9.57MY with a 95% confidence interval of 5.4-16.7My. Divergence time between C and D was estimated as 6.96My with a 95% confidence interval of 3.5 - 11.0My (Fig. 2).

### Species delimitation tests

Both Bayes factor delimitation (BFD) and Bayesian multi-species coalescent delimitation (BPP) analyses decisively supported the idea that *Dromiciops* is more than one species. In particular, the BFD marginal likelihood estimations were consistent and nearly identical for both the PS and SS approaches that included mtDNA and nDNA, supporting the idea of the existence of two *Dromiciops* species (i.e., clades A+ (BCD)) (2lnBf = -80.22 – - 4255 for PS and -70 – -4242 for SS) over alternative scenarios (Table S4, Fig 3a). However, the Bayesian multi-species coalescent delimitation analysis for the molecular data using BPP with A10 to determine a guide tree revealed a strong support for elevating the three main clades to the species level (A+ B + (C,D)) (posterior probability - pp=1).

**Figure 3.**
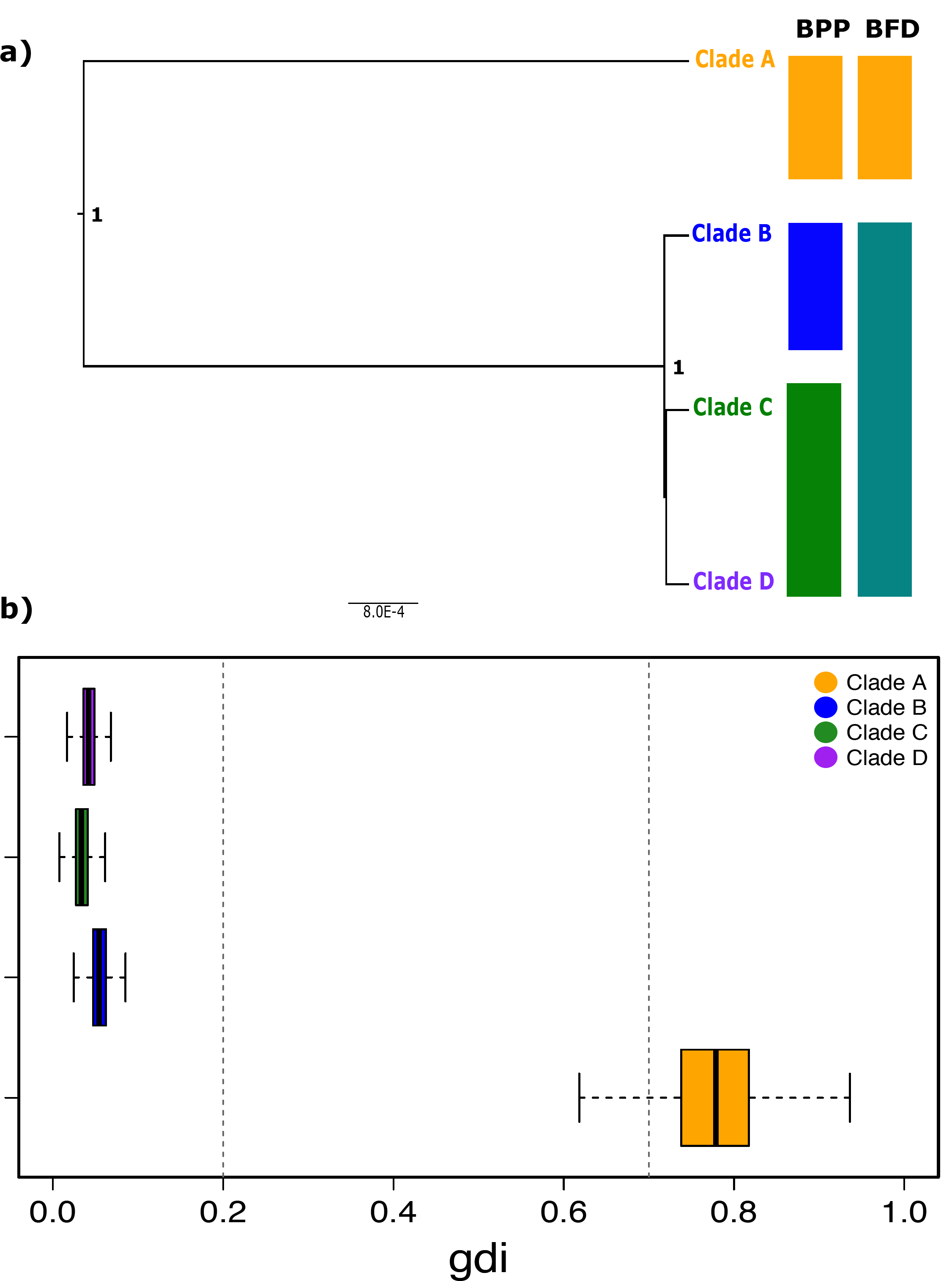
**a)** Schematic representation of the species delimitation hypotheses inferred by BPP using concatenated genes. Hypothesis were set as columns and colors denote clades. Numbers at each node of the tree indicate the posterior probability for that node inferred by analyses BPP. Color blocks identify distinct candidate species based in clades. Individual clades are same concatenated result of phylogeny. Clade A (Individuals from localities1-9), Clade B (Individuals from localities 10-12), Clade C (Individuals from localities 13-18) and Clade D (Individuals from localities 19-31) (see Table 1). **b)** Genealogical divergence index (*gdi*) for clades of *Dromiciops*. The dashed vertical bars indicate *gdi* thresholds. *gdi* values above 0.7 suggest separate species; gdi values below 0.2 are below the lower threshold for species delimitation; 0.2 < *gdi* < 0.7 are considered “ambiguous” range (Jackson et al. 2017).

Our results indicate that the best demographic model is that with small populations sizes (θ) and shallow divergence (τ) (i.e., prior scheme 3) (Fig S2; Table S5). This model proposes a moderate posterior probability (pp=0.8) for clade B and high posterior probability for all other nodes (pp=1). While clades B, C and D displayed gdi scores consistent with populations of a single species (*gdi* > 0.2) (Fig 3b), clade A (main: 0.77, 95%HPD 0.52-0.97) (Fig 3b), suggesting different species.

### Haplotype network analysis

Haplotype networks differed among genetic markers but three out of six displayed high concordance in their topology. These were the haplotype MJ networks for both mtDNA markers (Cytb and 12S RNA) and the nuclear gene IRBP (Fig S3). The Cytb haplotype network showed three clearly differentiated haplogroups (Fig S3a), where one group corresponds to samples from localities (1 to 9) north of 39°S. This group was separated by 13 mutational steps from the second group which is conformed only by four haplotypes and corresponds to samples from San Martin and Oncol. This second haplogroup was separated in turn by 9 mutational steps from a third group composed by only two haplogroups corresponding to samples from the southern localities (39.5-42°S; localities 13 to 30 in Fig S3). The haplotype network for 12s RNA was relatively similar to Cytb with the largest difference being the position of the haplotypes corresponding to Oncol and San Martin. The IRBP haplotype network differentiated populations from the north, but central and southern localities displayed shared haplotypes (Fig S3d).

### Population genetic structure

The amount of genetic variation explained by differences among groups was higher when samples where ordered in three groups (clade A, clade B, and clades C-D, Table 3) than when ordered in two groups (clade A vs the others). However, the amount of genetic variation explained by differences among localities within groups was systematically higher than that explained by differences among groups. The amount of genetic variation explained by differences among localities within groups was similar for both mtDNA and nDNA, but the amount of genetic differences among groups was higher for nDNA compared to mtDNA.

**Table 3.**
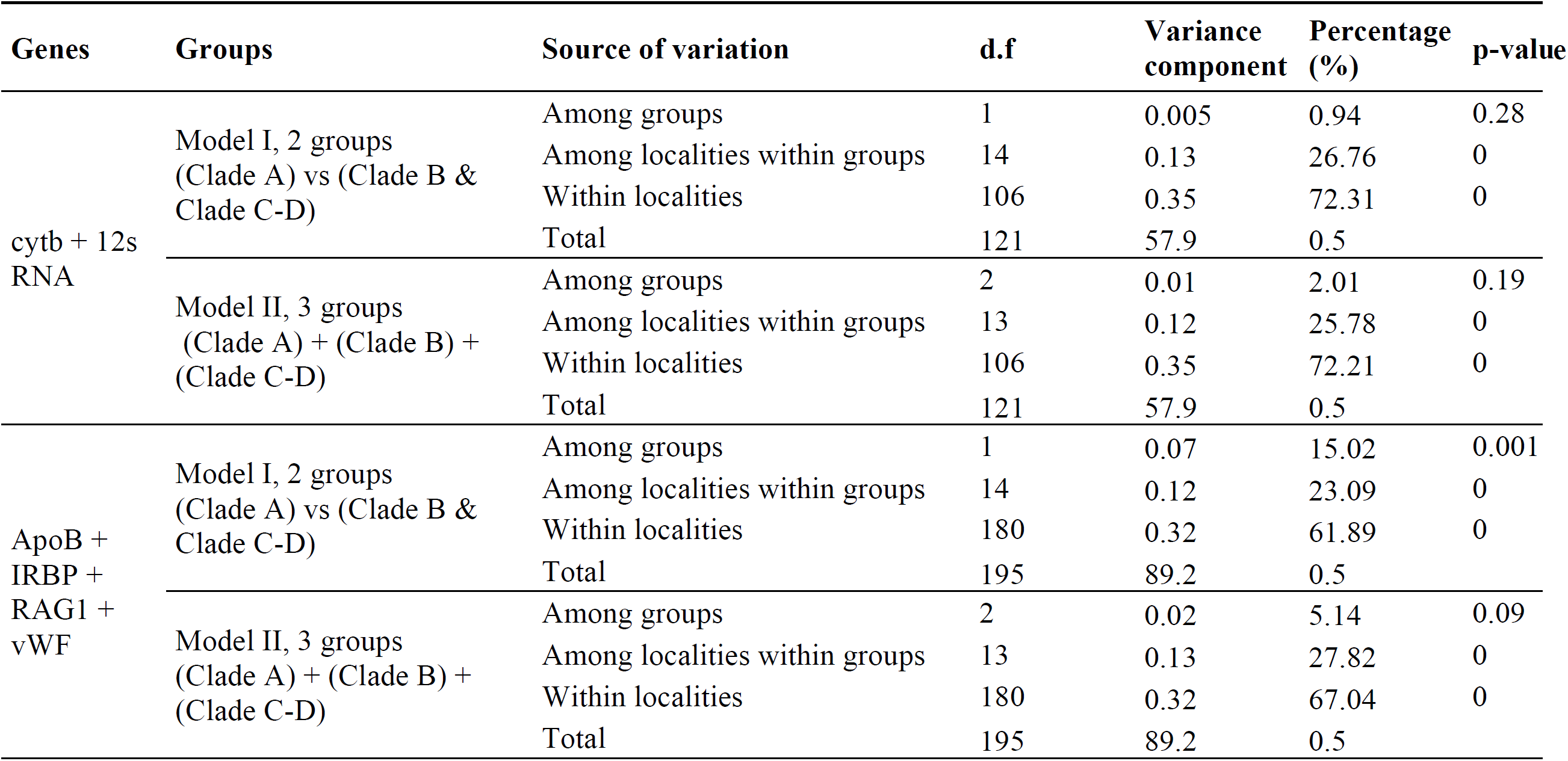
Analysis of Molecular variance result for mtDNA (Cytb + 12S RNA) and nDNA (ApoB + IRBP + RAG1 + vWF).

| Genes | Groups | Source of variation | d.f | Variance component | Percentage (%) | p-value |
| --- | --- | --- | --- | --- | --- | --- |
| cytb + 12s RNA | Model I, 2 groups (Clade A) vs (Clade B & Clade C-D) | Among groups | 1 | 0.005 | 0.94 | 0.28 |
|  |  | Among localities within groups | 14 | 0.13 | 26.76 | 0 |
|  |  | Within localities | 106 | 0.35 | 72.31 | 0 |
|  |  | Total | 121 | 57.9 | 0.5 |  |
|  | Model II, 3 groups (Clade A) + (Clade B) + (Clade C-D) | Among groups | 2 | 0.01 | 2.01 | 0.19 |
|  |  | Among localities within groups | 13 | 0.12 | 25.78 | 0 |
|  |  | Within localities | 106 | 0.35 | 72.21 | 0 |
|  |  | Total | 121 | 57.9 | 0.5 |  |
| ApoB + IRBP + RAG1 + vWF | Model I, 2 groups (Clade A) vs (Clade B & Clade C-D) | Among groups | 1 | 0.07 | 15.02 | 0.001 |
|  |  | Among localities within groups | 14 | 0.12 | 23.09 | 0 |
|  |  | Within localities | 180 | 0.32 | 61.89 | 0 |
|  |  | Total | 195 | 89.2 | 0.5 |  |
|  | Model II, 3 groups (Clade A) + (Clade B) + (Clade C-D) | Among groups | 2 | 0.02 | 5.14 | 0.09 |
|  |  | Among localities within groups | 13 | 0.13 | 27.82 | 0 |
|  |  | Within localities | 180 | 0.32 | 67.04 | 0 |
|  |  | Total | 195 | 89.2 | 0.5 |  |

The mantel test comparing geographical and genetic distances (Φ_ST_) using concatenated mtDNA markers and nDNA markers indicated a significant correlation between geographic and genetic distance matrices when considering all pairwise comparisons (mtDNA R^2^=0.15, p<0.01, R_XY_=0.36; and nDNA R^2^=0.31, p<0.01, R_XY_=0.56; Fig 4a-b). In particular pairwise differences that involved one locality from clade A vs localities from the other two clades displayed the highest Φ_ST_ for both mtDNA and nDNA (Fig 4c-d).

**Figure 4.**
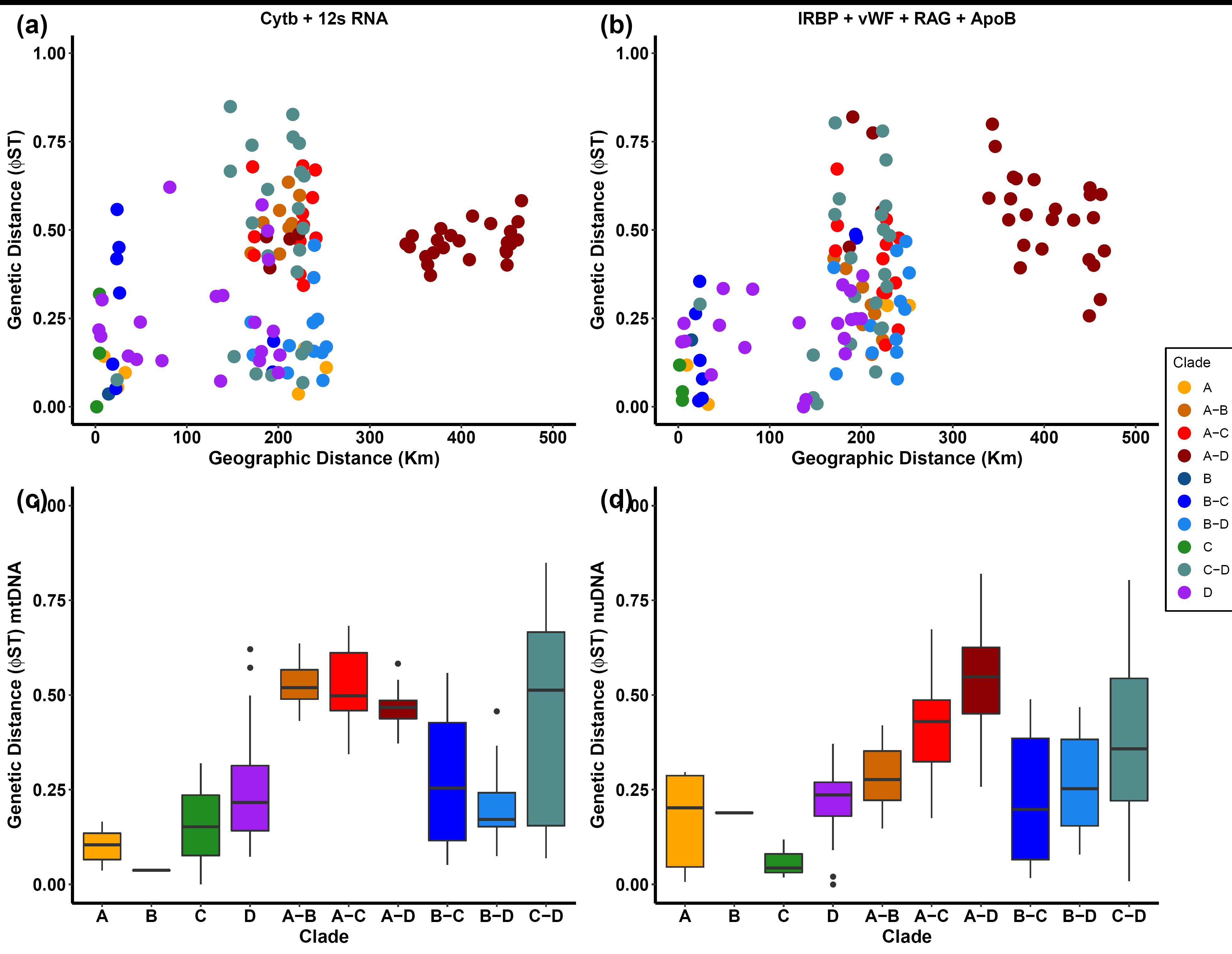
Pairwise geographic and genetic distances between all individuals for **a)** mtDNA (Cytb + 12S RNA) and **b)** nDNA (ApoB + IRBP + RAG1 + vWF). Comparation of genetic differentiation indexes (Φ_ST_) between clades and inter-clades for **c)** mtDNA and **d)** nDNA concatenated.

### Historical demography

Both Cytb Fu’s Fs and Tajima’s D statistics were negative for clades B, C and D but only statistically significant for clades B and D (Table 4). In addition, the mismatch distribution for Cytb gene indicated a unimodal distribution for clades B (Tau = 0.65), C (Tau = 1.04) and D (Tau = 0.34) but not for clade A, for which a multimodal mismatch distribution was fitted (Tau = 1.54) (Fig S4). Overall, these mtDNA results suggest that southern clades (B-C and D) but not A have undergone recent demographic expansions. In contrast, most nDNA genes had values not significantly different from zero.

**Table 4.** Neutrality tests for different clades. Fs= Fs’ Fu, D = Tajima’s D. Statistically significant values (p < 0.05) are indicated in bold

| Clade | Cytb |  | ApoB |  | IRBP |  | RAG |  | vWF |  |
| --- | --- | --- | --- | --- | --- | --- | --- | --- | --- | --- |
| | $F_s$ | $D$ | $F_s$ | $D$ | $F_s$ | $D$ | $F_s$ | $D$ | $F_s$ | $D$ |
| Clade A | 0.627 | 0.752 | 0.53 | -0.888 | 1.108 | -0.621 | 0.411 | -0.063 | 1.126 | -0.410 |
| Clade B | **-3.789** | **-2.025** | **1.969** | -0.376 | 0.081 | -0.323 | -0.306 | -0.552 | 1.004 | 0.518 |
| Clade C | -2,281 | -1.507 | **1.563** | -0.913 | -1.500 | -1.773 | 0.001 | 0.155 | 1.093 | 1.100 |
| Clade D | **-3.755** | **-1.844** | **2.096** | -0.273 | 0.969 | -0.779 | 0.491 | -0.177 | 0.529 | -0.889 |
| **Total** | -0.294 | 0.52 | **2.419** | -1.104 | -0.457 | -1.349 | 0.152 | 0.150 | 1.384 | -0.795 |

## Discussion

This study presents the most complete species-level genetic differentiation and phylogeographic assessment of one of the most intriguing relict marsupials, *Dromiciops*. By increasing the number of samples per locality and genes sequenced, we recovered the previously reported structure (Himes *et al.*, 2008), but also identified a novel, well-differentiated clade in the south. Likewise, our fossil-calibrated phylogeny suggests deep divergence times that go back to ∼13 Mya between clade A and the common ancestor of clades (B,C,D). This divergence would have occurred a few million years after the disappearance of other microbiotherids from the fossil record, in the Miocene (Goin & Abello, 2013). Thus, we conclude that clade A (northern localities) is a different species (*Dromiciops bozinovici*, as proposed by D’Elia et al. 2016). The other three clades have diversified more recently but still within the Miocene timescale, which is in sharp contrast with the previously hypothesized Pleistocenic scenario (Himes et al., 2008; see Fig 5).

**Figure 5.**
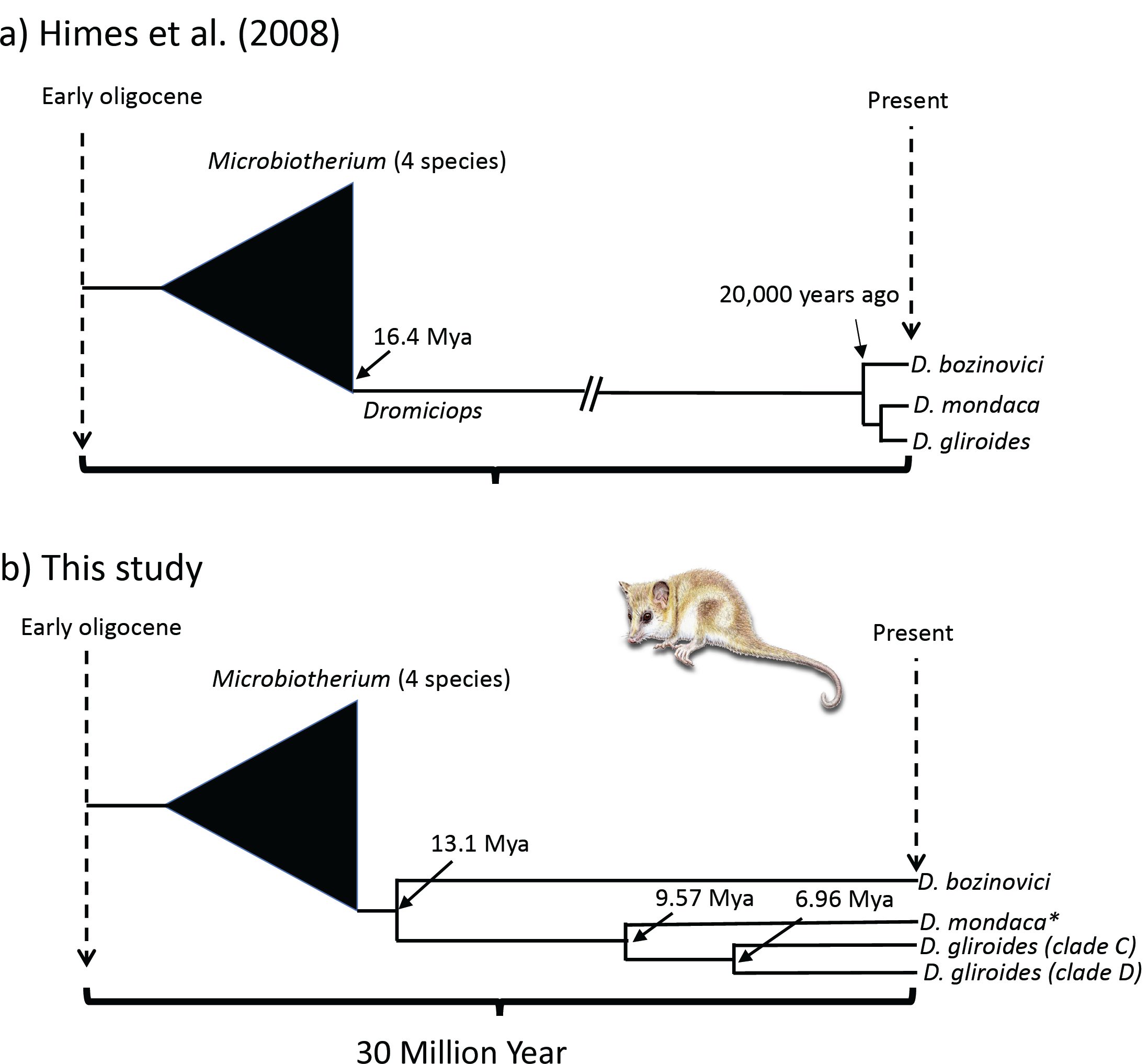
A schematic calibrated phylogeny depicting the conclusions of this study (clades are named according to D’Elia et al. 2016 propositions), in contrast with (**a**) the previously proposed model by Himes et al. (2008) who attributed the divergence of *Dromiciops* clade to recent Pleistocenic events. In this study (**b**), the calibrated divergence times indicate an early speciation event producing *D. bozinovici* soon after the disappearing Microbiotherium species from the fossil record (Goin and Abello, 2013). Asterisk denote a putative species as proposed by D’Elia et al. (2016) which however did not surpass the *gdi* threshold. Branch lengths and splits are scaled to the whole period of 30 My.

Our estimated percentage of divergence in mtDNA between clade A (*D. bozinovici*, hereafter) and the other clades (range: 7.3-12%) is comparable with what Himes et al. (2008) reported (range: 11.3-15%). While these authors did not attribute these divergences to a major split in the *Dromiciops* lineage, these levels of divergence are typical for inter-specific variation in mammals (Baker & Bradley, 2006; Schrago & Mello, 2020), and led D’Elia et al. (2016) to revisit the problem and designing clades A and B to the level of species (Table 5). In contrast, Suárez-Villota *et al.* (2018), analyzing four nuclear and two mtDNA markers in *Dromiciops* individuals from 20 localities reported genetic distances (K2P) averaged across all loci that were an order of magnitude lower (on the range of 0.001 to 0.015%) and concluded that these differences were within the intra-species range previously reported for other marsupials. Unfortunately, the reported genetic distances indicated by Suarez-Villota et al., 2018 are only comparable if the same genes are used, as substitution rates can vary greatly among genes (an order of magnitude, between nuclear and mitochondrial genes, in our case). Thus, it is likely that their K2P estimations are low because they included highly conserved nuclear genes in the estimations. Finally, a recent and extensive compilation of more than 57000 mammal cytb sequences including 82 marsupial species showed that 95% of cytb interspecies genetic distances in marsupials fall between 9% and 16% with a mean of 12.3% (Schrago & Mello, 2020). Divergence between D. *bozinovici* and the other *Dromiciops* clades falls well within this range.

**Table 5.** Summary of studies of clinal population variation in *Dromiciops.*

| Reference | Geographic coverage (individuals / localities) | Type of evidence | Method | Main conclusion | Proposed scenario according to Fig 6 |
| --- | --- | --- | --- | --- | --- |
| (Himes et al. 2008) | 56 / 21 | 2 mtDNA genes | standard population genetics | Deep phylogeographic structure | “Pleistocene” |
| (D'Elia et al. 2016) | 56 / 10 | morphological | geometrics morphometrics and qualitative analysis | Discrete groups with new species proposition | none |
| (Valladares-Gómez et al. 2017) | 69 / 16 | morphological | geometrics morphometrics | Clinal differentiation | “Pleistocene” |
| (Suarez-Villota et al. 2018) | 45 / 20 | 4 Nuclear and 2 mtDNA partial sequences | species delimitation & coalescent analysis | Absence of differentiation | none |
| (Martin 2018) | 41 / 30 | morphological | classic morphometrics | Absence of clinal differentiation | “Pleistocene” |
| (Valladares-Gomez et al. 2019) | 47 / 9 | 11 microsatellites | standard population genetics | Existence of discrete groups | “Pleistocene” |
| This study | 159 / 31 | 2 mtDNA genes and 4 nuclear genes | standard population genetics & coalescent analysis | Deep phylogeographic structure confirming *D. bozinovici* as different species | “Miocene” |

Fossil calibrations suggested a particularly old crown age for *Dromiciops* (mean of 13.1 My, middle Miocene), whose diversification would have happened soon after the separation of *Dromiciops* from the extinct genus *Microbiotherium*, in the early Miocene (Goin *et al.*, 2007; Goin & Abello, 2013). These estimations, however, should be viewed with caution, as our credible intervals are relatively large (95% CI: 6.4 - 25.3 My). First, our data set has a greater number of individuals in the ingroup than in the outgroup, an unbalance that could affect molecular clock estimations (i.e., influence of taxon sampling) (Rosenberg & Kumar, 2003; Nabhan & Sarkar, 2012; Soares & Schrago, 2012). Second, our phylogenetic reconstruction was performed with incomplete gene matrices (not all genes were available for all individuals) increasing the uncertainty of divergence rates. Third, we used fossils from marsupials of South America (characterized by old divergence times: 10-35MY) and Australia (more recent: 3-15MY) for calibration, whose varying range could affect confidence intervals (Mitchell *et al.*, 2014; Eldridge *et al.*, 2019). Given the position of *Dromiciops* in the marsupial phylogenetic tree (Nilsson *et al.*, 2004; Mitchell *et al.*, 2014), the introduced uncertainty due to the fossil outgroup designation could inflate credible intervals of the ingroup (Herrera & Dávalos, 2016; Lee, 2016). Still, our three estimated divergences fall within the Miocene, which is coincident with episodic marine transgressions that isolated large portions of land during this period.

The middle Miocene transgression (MMT) was a massive flooding of seawater that invaded the western face of Los Andes between 38-48° S (Malumián & Náñez, 2011), splitting Chile into an extensive Northern territory, which could have isolated a large population of *Dromiciops* for enough time to differentiate into *D. bozinovici* (see Fig 5). According to Malumián and Náñez (2011), two portions of land were not covered by the LMT and survived the flooding as islands: the coastal range near Valdivia (where clade “B” of *Dromiciops* is currently found) and a southern part of Chiloé island, which would have been the origin of our clade “C” and “D”(see Fig 6). Interestingly, paleolandscape reconstructions of the boundary basins left by the MMT during the Miocene and the Pliocene are coincident with the proposed ancestral distribution of *Nothofagus* lineages in Chile and Argentina (Acosta *et al.*, 2014). Moreover, these authors show that most of the divergence in *Nothofagus* took place during the Miocene, as in *Dromiciops*. Then, the tendency Microbiotherids to evolve in close coevolutionary association with the so-called *Nothofagus*-*Chusquea* biome (Hershkovitz, 1999: pp. 7-8), together with the strong diversifying force of the marine transgressions, likely acted as diversifying force in the region (Acosta *et al.*, 2014). This is coincident with our results fiving to Clade A (*D. bozinovici*) an older history of expansion, as these populations were outside the boundary basins left by marine transgressions, whereas clades B and C would have originated from highland refuges south of the Arauco Basin (Fig 6a). Our results showing commonalities between Chiloé Island, mainland and Argentina (clade D) would be explained by the fact that (1) the South of the island survived to the floodings and (2) the island was probably connected to mainland during the last 1 – 1.5 Mya (Acosta *et al.*, 2014).

**Figure 6.**
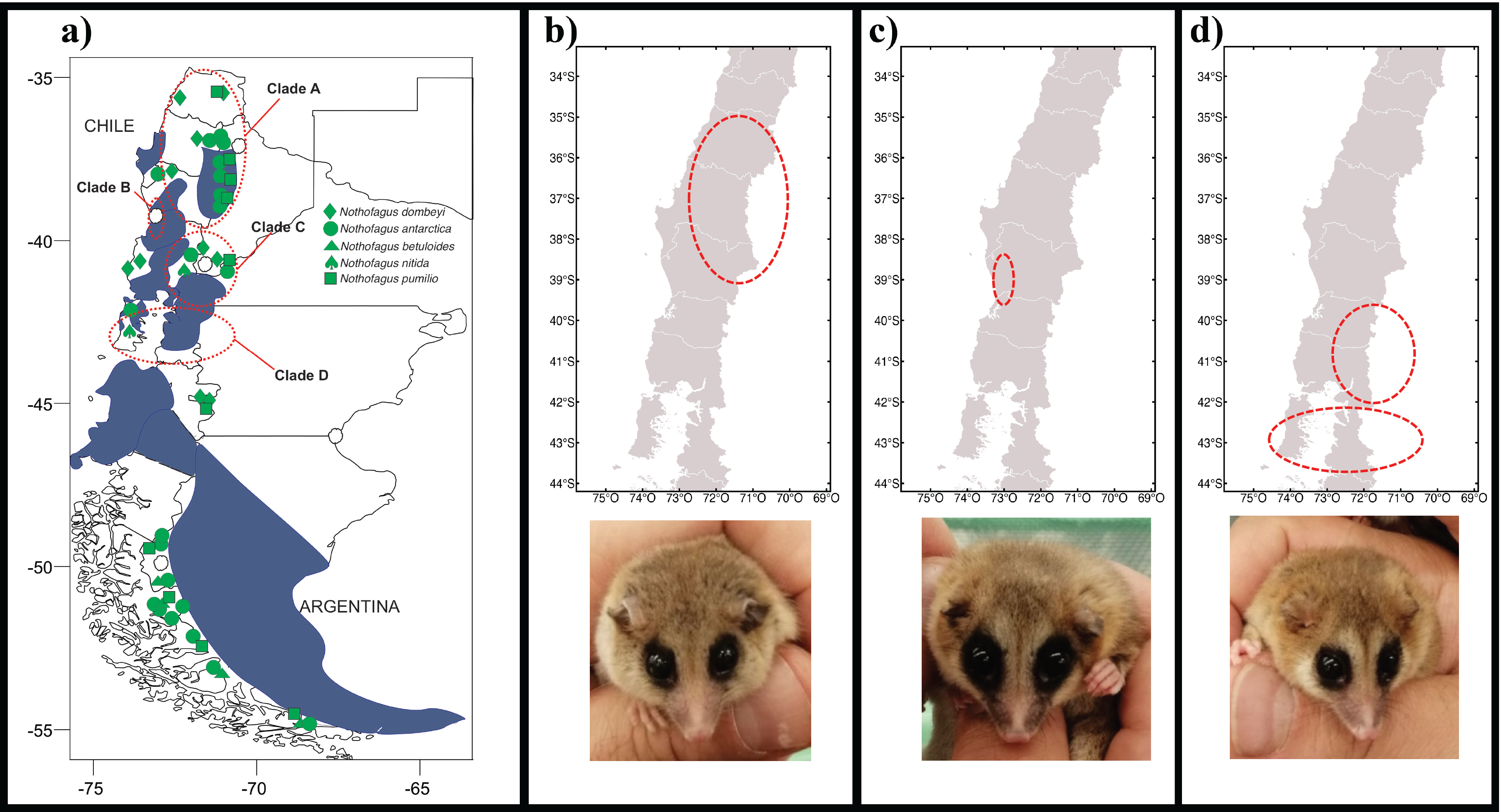
**a)** Paleolandscape reconstruction of southern South America (modified from Acosta et al., 2014) showing the Miocene basins (blue), the hypothetical distribution of *Nothofagus* lineages (green), and the approximate distribution of *Dromiciops* clades found in this study (red ovals). White areas within basins indicate mountain that became islands when marine transgression occurred. In **b) – d)** a detail of the distribution together with live specimens of each clade are shown (photo: R. Nespolo).

Summing up, our conclusions adds to a paradigm shift for retracing the evolutionary history for Southern South America to a new and older geo-climatic context, which contrasts with the more recent quaternary glaciations scenario that were previously accepted (see a Discussion in Acosta et al., 2014). Further studies are warranted, in order to determine the conservation status of the new *D. bozinivici* species at the northern range, as well as the isolation of the southern *Dromiciops* clades in their relationship with *Nothofagus* forests that distributes over the Patagonia.

## Acknowledgments

This study was funded by FONDECYT grant No 1180917 (R.F. Nespolo) and ANID PIA/BASAL FB0002. Partial funding was also provided by DID grant Universidad Austral de Chile S 2018-20 (P Saenz-Agudelo). CONICYT PhD fellowship 2016 N° 21160901 (J.F Quintero-Galvis). We appreciate the insightful suggestions of Guillermo D’Elia and Francisco Fontúrbel to the draft.

